# Proteomic reprogramming of ileal epithelial cells during homologous superimposed intestinal trematode infection reveals coordinated restoration of intestinal homeostasis

**DOI:** 10.64898/2026.04.29.720092

**Authors:** Emma Fiallos, Paola Cociancic, José Guillermo Esteban, Carla Muñoz-Antoli, Rafael Toledo

**Author notes:** Corresponding author: (RT).

## Abstract

**Background:** Intestinal helminth infections trigger complex host responses, determining parasite survival and tissue homeostasis. Primary *Echinostoma caproni* infection disrupts epithelial metabolism, differentiation, and repair in an IL-25-deficient environment, as shown in a previous study by our research group; however, the adaptive mechanisms during homologous superimposed infections remain unclear.

**Methodology/Principal findings:** Male ICR mice were assigned to control, primary infection, and homologous superimposed infection groups, and ileal epithelial cells were isolated for proteomic profiling using liquid chromatography–tandem mass spectrometry (LC–MS/MS) with data-dependent acquisition (DDA) and sequential window acquisition of all theoretical mass spectra (SWATH). Differential protein expression was analyzed with Elastic Net regression, partial least squares discriminant analysis, and fold-change ranking, while functional enrichment and protein–protein interaction networks were explored using gene set enrichment analysis (GSEA) and STRING. Notably, homologous superimposed infection revealed proteomic signatures associated with lysosomal and peroxisomal lipid metabolism, PPAR pathway activation, cytoskeletal reorganization, epithelial barrier reinforcement, a specialized antimicrobial peptide repertoire, and interactions between IgE receptor-associated proteins, consistent with a restoration of intestinal homeostasis influenced by IL-25.

**Conclusions:** Host adaptation to repeated *E. caproni* exposure involves coordinated metabolic, signaling, and tissue repair responses that partially restore intestinal homeostasis, with IL-25 emerging as a central regulator linking metabolic reprogramming, epithelial integrity, and anti-helminth immunity, thereby providing a proteomic framework for understanding how repeated helminth exposure drives partial resistance through integrated epithelial and immunometabolic adaptations.

**Author summary:** We investigated how repeated intestinal worm infections affect the cells lining the small intestine in mice. Infections with intestinal trematodes can disrupt the normal balance of the gut, leading to tissue damage and altered metabolism. Using a proteomics approach, we measured changes in thousands of proteins in intestinal epithelial cells during a first infection and after a second, repeated infection. We found that the first infection caused stress in the cells, impaired oxygen use, and reduced the activity of pathways that normally help repair tissue. In contrast, the repeated infection triggered a coordinated response that restored many cellular functions. Cells increased protein activity related to fat metabolism, tissue structure, barrier integrity, and antimicrobial defense. We also observed evidence that the immune signaling molecule interleukin-25 plays a central role in coordinating these protective and repair processes. These results suggest that the gut epithelium can adapt to repeated infections by reorganizing its metabolic and structural functions, which may help limit tissue damage and promote partial resistance to parasites. Our study provides a detailed map of the molecular changes that underlie this adaptation, improving our understanding of how the intestinal lining responds to repeated worm infections.

## Introduction

Intestinal helminth infections remain a major public health concern in many regions of the world, affecting millions of people and animals through chronic morbidity, malnutrition, and impaired development affecting more than 1.5 billion people worldwide [1–4]. These parasites have evolved sophisticated strategies to evade or modulate host immune responses, often establishing long-term infections in the gastrointestinal tract. Among the key host factors determining susceptibility or resistance to intestinal helminths, interleukin-25 (IL-25, also known as IL-17E) has emerged as a central alarmin cytokine produced predominantly by intestinal epithelial cells, particularly tuft cells, in response to helminth-derived signals. IL-25 initiates a type 2 immune cascade by activating innate lymphoid cells type 2 (ILC2s) and promoting Th2 polarization, which drives effector mechanisms such as goblet cell hyperplasia, mucus hypersecretion, antimicrobial peptide production, smooth muscle hypercontractility, and epithelial barrier reinforcement, collectively facilitating parasite expulsion [5,6].

The *Echinostoma caproni*–ICR mouse model has proven to be a valuable experimental system for investigating the immunological and pathophysiological mechanisms underlying susceptibility and acquired resistance to intestinal trematodes. In primary infections, ICR mice are highly susceptible, supporting long-lasting infections with high worm burdens and minimal expulsion. This susceptibility is associated with a deficient IL-25 response, leading to disrupted epithelial metabolism, impaired differentiation and repair, sustained oxidative stress, and failure to mount effective type 2 immunity. In contrast, mice that clear a primary infection (e.g., via praziquantel treatment) or are exposed to repeated homologous infections develop partial resistance, characterized by a marked reduction in worm establishment and survival upon challenge [7–9]. Recent studies have demonstrated that this “concomitant immunity” or partial resistance to superimposed homologous *E. caproni* infections is largely dependent on elevated IL-25 levels rather than classical Th2 responses, as blockade of IL-25 abrogates protection while interference with downstream Th2 effectors (e.g., IL-4Rα or IL-13Rα2) does not fully prevent resistance [8,10].

Despite these advances, the adaptive mechanisms at the epithelial level during repeated exposures remain poorly characterized at the molecular level. Previous proteomic analyses from our group revealed that IL-25-mediated resistance in this model involves profound reprogramming of ileal epithelial cells, including restoration of metabolic homeostasis, enhancement of barrier integrity, and modulation of inflammatory pathways. However, these studies focused primarily on primary infections or IL-25-treated scenarios, leaving unexplored the proteomic landscape during homologous superimposed infections, where partial resistance develops naturally in the presence of ongoing or recent IL-25-driven responses [5,6,11].

In the present study, we hypothesized that repeated exposure to *E. caproni* induces coordinated proteomic adaptations in ileal epithelial cells that support partial restoration of intestinal homeostasis and contribute to the observed resistance, with IL-25 acting as a key orchestrator linking immunometabolic reprogramming, cytoskeletal reorganization, epithelial barrier reinforcement, and specialized antimicrobial defenses. To test this, we performed label-free quantitative proteomics (DDA + SWATH) on isolated ileal epithelial cells from ICR mice subjected to primary infection or homologous superimposed infection, compared with uninfected controls. Differential expression was analyzed using multivariate approaches (penalized multiple regression models), followed by pathway enrichment Gene Set Enrichment Analysis (GSEA) with Gene Ontology (GO) and Kyoto Encyclopedia of Genes and Genomes (KEGG), and protein–protein interaction networks (STRING) to uncover global functional reprogramming beyond individual protein changes.

These findings provide the first proteomic framework for understanding how repeated helminth exposure drives partial resistance through integrated epithelial and immunometabolic adaptations, highlighting IL-25 as a pivotal regulator and offering insights into potential targets for modulating host responses in helminthiasis.

## Material and methods

### Animals and infection procedures

An experimental study was conducted using the *Echinostoma caproni*–rodent model. The *E. caproni* intestinal trematode strain and infection procedures have been previously described [12]. Encysted metacercariae of *E. caproni* were obtained by microdissection of kidneys and pericardical cavities of experimentally infected *Biomphalaria glabrata* snails. Subsequently, 12 male ICR (CD1) mice (*Mus musculus*), weighing 30–35 g, were randomly assigned to three experimental groups (n=4 per group): non-infected control, primary infection, and homologous superimposed infection. Experimental infection was performed by gastric gavage orally by administering 25 *E. caproni* metacercariae per mouse using a gastric cannula, following established protocols [13]. Homologous superimposed infection was induced at 2 weeks post-primary infection (wpi). Animals were euthanised at 4 and 8 wpi, and necropsy was performed for adult worm recovery and intestinal tissue collection. This time point was chosen because it corresponds to parasite maturation, coincides with the expulsion of worms in resistant hosts, and enables direct comparison with previous studies. This fact does not discard that additional changes can be observed at other time points.

### Quantification of IL-25 expression by real-time PCR

To access IL-25 expression in the ileum, total RNA was extracted from full-thickness ileum tissue collected from necropsied mice. RNA isolation was performed using Realclean Tissue/Cells RNA kit (Real Laboratory) following the manufacturer’s protocol. The cDNA was synthesized from the extracted RNA using the High Capacity cDNA Reverse Transcription kit (Applied Biosystems). Quantitative real-time PCR was carried out using a TaqMan^®^ Gene Expression Assay specific for mouse IL-25 (Mm00499822_m1, Applied Biosystems). For each reaction, 40 ng of RNA-derived cDNA was combined with 10µL of TaqMan^®^ Universal PCR Master Mix (2x, No AmpErase^®^ UNG), 1µL of the TaqMan^®^ assay, and nuclease-free water to reach a final volume of 20µL. Amplification reactions were performed in 96-well plates using an Abi Prism 7000 (Applied Biosystems). Thermal cycling conditions consisted of an initial denaturation setup of 10 min at 95 °C, followed by 40 cycles of 15 s at 95 °C and 1 min of annealing/extension at 60 °C each. All samples, endogenous control, and negative controls were analyzed in triplicate. Each assay included two unlabeled primers, and a 6-FAM™ dye-labeled, TaqMan^®^ MGB probe.

Gene expression levels were determined by calculation the cycle threshold (Ct) of each sample. To correct for variations in RNA quantity and reverse transcription efficiency, expression levels were normalized to the housekeeping gene β-actin. Relative IL-25 expression was calculated using the comparative method 2^-ΔΔCT^ method, with values standardized those obtained from non-infected control mice.

### Parasite recovery and protein extraction

The small intestine was removed at necropsy, opened longitudinally, and adult worms were recovered and counted under a stereomicroscope. For proteomic analysis, intestinal epithelial cells were isolated as previously described by Muñoz-Antoli et al. (2014) [14]. Protein extracts were prepared from intestinal epithelial cells. Intestinal tissue was gently rinsed with ice-cold Hank’s Balance Salt Solution supplemented with 2% heat-inactivated fetal calf serum. The supernatant was discarded and replaced with fresh washing buffer; this procedure was repeated at least 4 times until the wash solution remained clear. The tissue was then cut into approximately 1 cm fragments and incubated for 20 min at 37 °C in a dissociation buffer consisting of HBSS supplemented with 10% fetal calf serum, 1 nM EDTA, 1 mM DTT, 100 U/ml penicillin, and 100 μg/ml streptomycin. The supernatant was collected and kept on ice, and the incubation was repeated using fresh buffer. Supernatants were pooled and filtered through a 100 µm cell strainer. Intestinal epithelial cells were then pelleted by centrifugation at 200 g for 10 min at 4°C and washed 3 times with PBS under identical conditions to remove residual reagents.

For protein extraction, processed tissue (1 g) was transferred to a 50 mL Falcon tube containing 15 mL of T-PER reagent, supplemented with 2.5 mL of protease inhibitor cocktail (ethylenediaminetetraacetic acid or EDTA, Roche) and 17.5 µl of phosphatase inhibitor (Sigma Aldrich), to final concentrations of 1X and 0.1%, respectively. Tissue homogenization was achieved by 3 cycles of alternating vigorous vortexing and vertical agitation, each lasting 5 min. Homogenates were subsequently transferred to 15 mL tubes and centrifuged twice at 5000 g for 5 min at 4 °C to clarify the extracts. The resulting supernatants were aliquoted into 1.5 mL microcentrifuge tubes and stored until further analysis.

### LC–MS/MS proteomic analysis

Differential protein expression was analysed by liquid chromatography–tandem mass spectrometry (LC–MS/MS) using data-dependent acquisition (DDA) and sequential window acquisition of all theoretical mass spectra (SWATH) at the Proteomics Unit of the SCSIE (University of Valencia).

For spectral library generation, representative protein extracts from each experimental group were pooled, separated by SDS-PAGE, and subjected to in-gel enzymatic digestion with sequencing-grade trypsin (Promega). Samples were incubated with 500 ng trypsin in ammonium bicarbonate at 37 °C; digestion was stopped with 10% trifluoroacetic acid (TFA), and peptides were extracted with pure acetonitrile (ACN), dried under vacuum, and resuspended in 20 µl of a 2% ACN containing 0.1% TFA.

Samples were randomised prior to analysis to minimise batch effects. For LC-MS/DDA, 5 µl of peptides were injected onto a trap column (3 µm C18-CL, 120 Å, 350 µm × 0.5 mm; Eksigent), desalted with 0.1% TFA for 3 min at 5 µl/min, and separated on an analytical column (3 μm particle size C18-CL, 120 Ᾰ, 75 μm diameter × 150 mm; Eksigent). Peptides were eluted using a linear gradient of 7–40% ACN in 0.1% formic acid (FA) at 300 nl/min for 60 min. Survey MS1 scans were acquired over an m/z range of 350–1250 for 250 ms. MS2 scans were acquired in ‘high sensitivity’ mode over 100–1500 m/z for 150 ms, with quadrupole resolution set to ‘UNIT’. Charge of 2+ to 5+ and a minimum intensity of 70 counts per second were selected, with up to 25 ions for fragmentation after each survey scan. Dynamic exclusion was set to 15 s.

For SWATH analysis, 20 µg of protein per sample were processed by SDS-PAGE and digested as described above. Five microlitres of peptides were analysed under similar chromatographic conditions using a 7–37% ACN gradient in 0.1% FA at 300 nl/min. The TripleTOF instrument was operated in swath mode, in which a 0.050-s TOF MS scan from 350–1250 m/z was performed, followed by product ion scans of 0.080 s across 100 variable windows (400–1250 m/z). The total cycle time was 2.79 s. All analyses were performed on an Eksigent nanoLC 425 system coupled to a 6600plus TripleTOF spectrometer (ABSCIEX), operated in DDA or SWATH mode as appropriate.

### Protein identification and quantification

Protein identification was carried out using ProteinPilot v5.0 (ABSCIEX) with the Paragon algorithm [15], using trypsin specificity, iodoacetamide alkylation of cysteines, searching against the SwissProt database with and without taxonomic restriction to *M. musculus*. A false discovery rate (FDR) of 1% was applied at peptide and protein levels, and protein grouping was performed using the Pro Group algorithm.

SWATH-based protein quantification was performed using PeakView 2.2, employing endogenous peptides for retention time calibration to align retention times. Peptide extraction parameters were predefined, and protein areas were normalised to the total signal per sample. Data were exported for statistical analysis.

### Statistical, functional enrichment and network analyses

Adult worm burdens were compared using Student’s t-test, with significance set at p < 0.05. Expression levels of IL-25, as measured by PCR, were compared among groups using a one-way analysis of variance (ANOVA) with Bonferroni test as post-hoc analysis (p < 0.05). Data were log-transformed to achieve normality and analysed using GraphPad Prism. Differential protein expression was assessed using penalized multiple regression models (Elastic Net), principal component analysis (PCA), and supervised partial least squares discriminant analysis (PLS-DA). Differentially expressed proteins were identified using the *Limma* package in R, ranked by p-value and corrected using the Benjamini–Hochberg method; proteins with adjusted p ≤ 0.05 were selected (R script in S1 and S2 Files).

Functional interpretation was performed using gene set enrichment analysis (GSEA) to identify significantly enriched biological processes and pathways [16], using GO and KEGG databases as references. For GSEA-GO proteins were categorized into three GO domains: molecular function, biological processes and cellular components. Statistical significance was estimated by gene label permutations and adjusted using the Benjamini–Hochberg method. Results were expressed as normalised enrichment scores (NES), and the 20 most enriched terms were selected based on the highest NES values. All analyses were conducted using Bioconductor’s Clusterprofiler.

Protein–protein interaction networks were constructed using the STRING platform [17] based on proteins associated with the 20 most representative GO-based GSEA terms (10 upregulated and 10 downregulated). Networks were generated using default parameters, and functional modules were identified using k-means clustering.

### Ethics statement

The animals were maintained under conventional conditions with food and water *ad libitum*. This study has been approved by the Ethical Committee of Animal Welfare and Experimentation of the University of Valencia (Ref#A18348501775). Protocols adhered to Spanish (Real Decreto 53/2013) and European (2010/63/UE) regulations.

## Results

### Infection rates

During primary *E. caproni* infection, a high number of adult worms was recovered per mouse, ranging from 17 to 25 (22.0 ± 3.8), corresponding to an infection rate of 88%. Primary infection was characterized by a high parasite load, whereas mice subjected to homologous superimposed infection exhibited a markedly reduced infection rate of 5%, indicating the development of partial resistance to repeated exposure to metacercariae.

### IL-25 gene expression

Quantitative real-time PCR analysis revealed differential IL-25 expression among the experimental groups (Fig 1). Compared with non-infected controls, mice undergoing primary infection showed a slight increase in IL-25 transcription. In contrast, mice subjected to homologous superimposed infection displayed a markedly higher IL-25 expression. Statistical analysis indicated that IL-25 expression was significantly higher in the superimposed infection group compared with both the control and primary infection groups, whereas the increase observed during primary infection was modest. These results demonstrate a pronounced upregulation of IL-25 transcription associated with repeated exposure to *E. caproni*.

**Fig 1.**
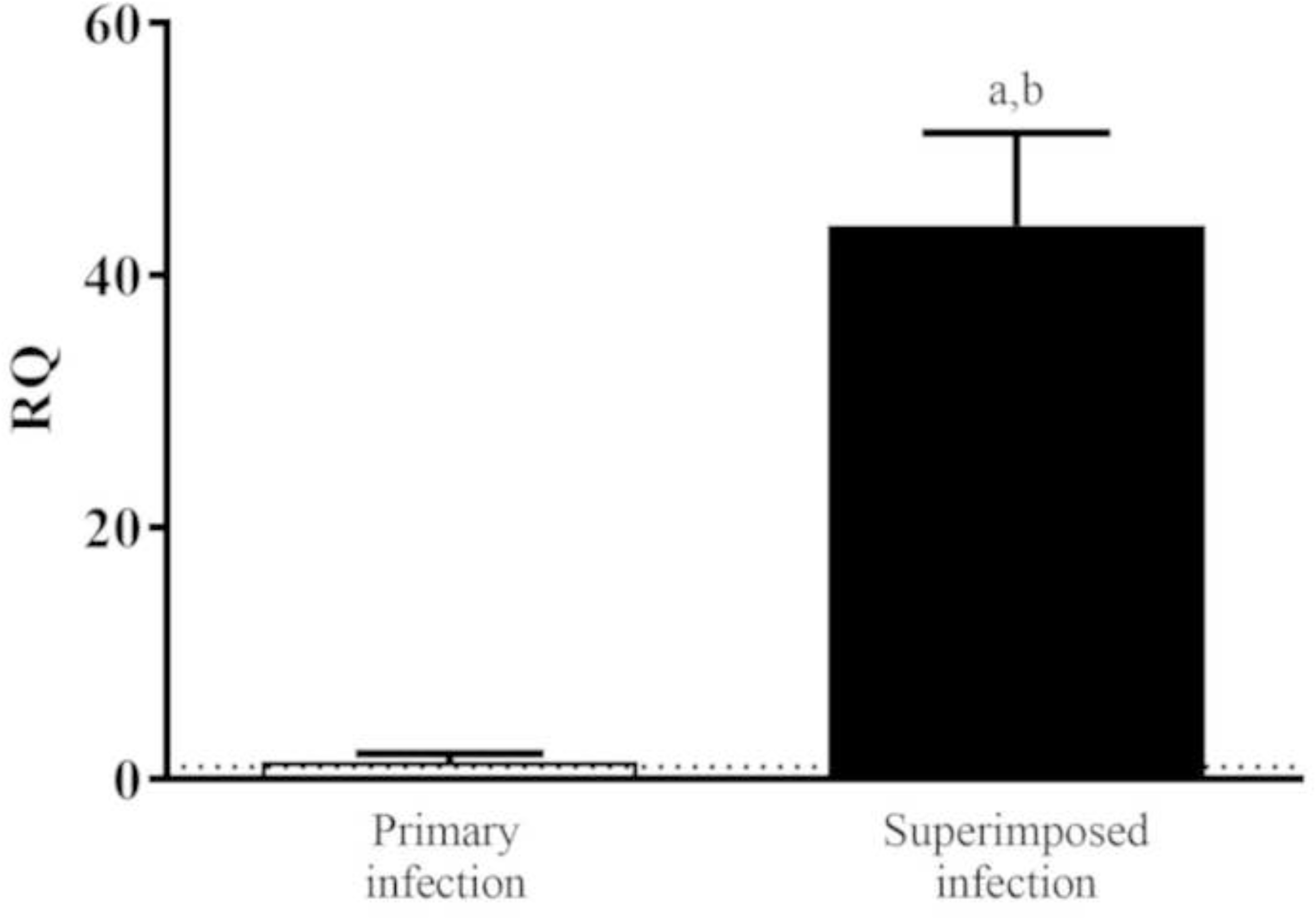
Interleuquin-25 expression in ileum of ICR mice during primary and superimposed *Echinostoma caproni* infections. Interleuquin-25 (IL-25) expression was quantified by real-time PCR, normalized to β-actin, and standardized to the relative amount against day 0 sample. Vertical bars represent standard deviation. Letters indicate significant differences: a, versus negative control; b, versus primary infection (p < 0.05).

### Identification of differentially expressed proteins

A total of 2,196 proteins were identified in the spectral library at a 1% FDR, and 2,281 proteins were identified when the database was restricted to *Mus musculus* (FDR 1%). Of these, 1,809 proteins were quantified across the 12 samples corresponding to the three experimental groups, maintaining an FDR < 1%. To identify differentially expressed proteins, a regularized Elastic Net regression model was applied and optimized by cross-validation using the caret package in R (λ = 0.295 and α = 0.4). Results were visualized using hierarchical clustering dendrograms and heatmaps, revealing sample grouping according to experimental condition (Fig 2A). PCA explained 47% of the total variance (PC1: 36%, PC2: 12%) (Fig 2B), while PLS-DA confirmed the observed separation, with the first two components explaining 40% and 16% of the variance, respectively (Fig 2C). Protein selection based on the Elastic Net model combined with variable importance in projection (VIP > 1.5) from PLS-DA identified two main clusters, distinguishing proteomic profiles among experimental groups (Fig 2D).

**Fig 2.**
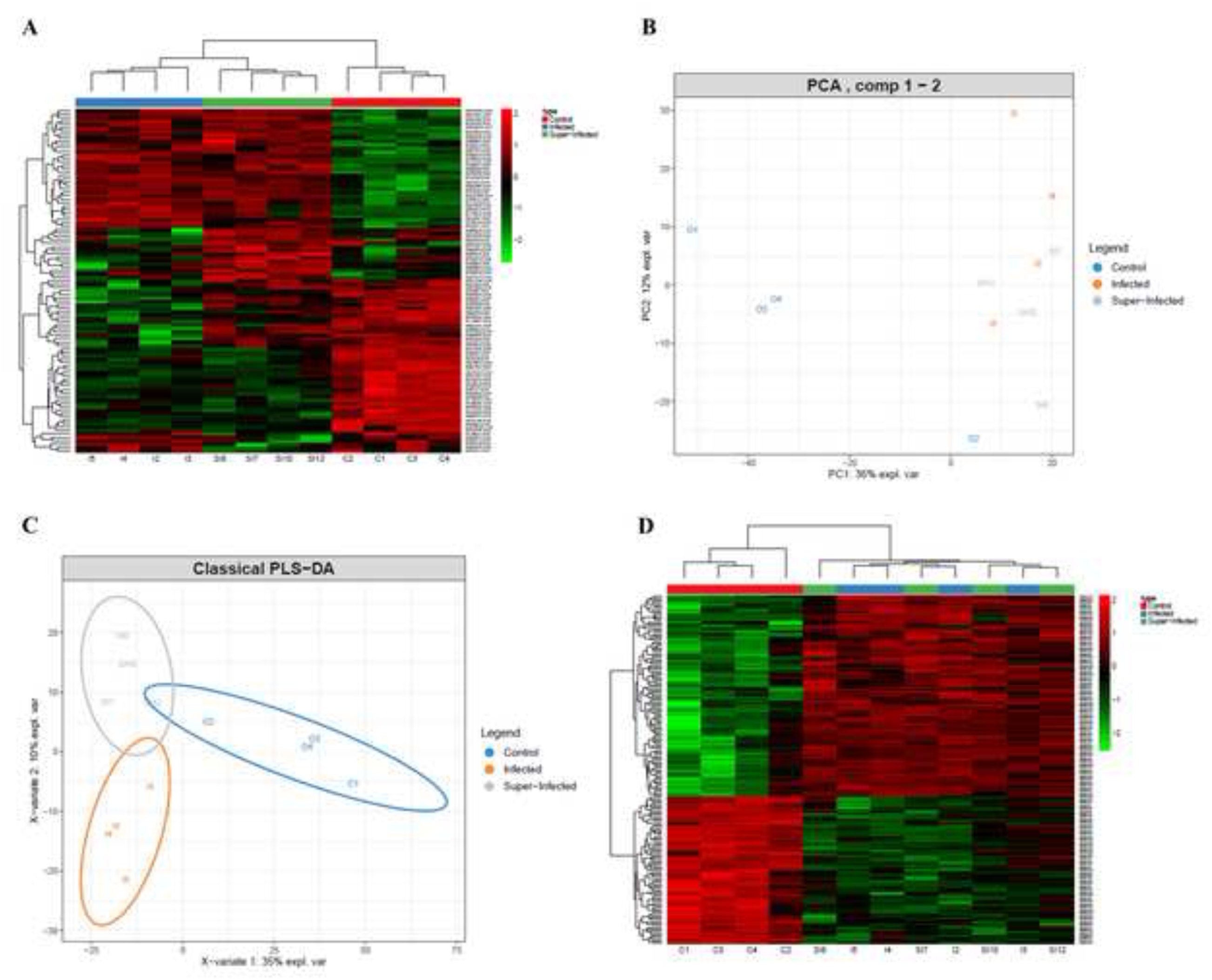
Differential protein expression in ileum of ICR mice during primary and superimposed *Echinostoma caproni* infections. Hierarchical heatmap generated from ElasticNet analysis, showing protein expression among the different conditions studied **(A)**. Proteins were analyzed by principal component analysis (PCA) **(B)** and supervised classification by partial least squares discriminant analysis (PLS-DA) **(C)**. These proteins are presented in the hierarchical heatmap with a VIP >1.5 in the PSL-DA classification **(D)**.

Overall, these results indicate that both primary and homologous superimposed infection with *E. caproni* significantly alter the intestinal proteome compared with the control group, whereas no significant individual protein-level differences were detected between primary infection and homologous superimposed infection. Nevertheless, the 1,809 quantified proteins were ranked according to fold change values obtained using the *Limma* model to identify coordinated alterations in metabolic pathways or signaling cascades through GSEA and STRING analysis.

Fig 3 presents a schematic overview of the workflow used for the identification of differentially expressed proteins under the experimental conditions analyzed.

**Fig 3.**
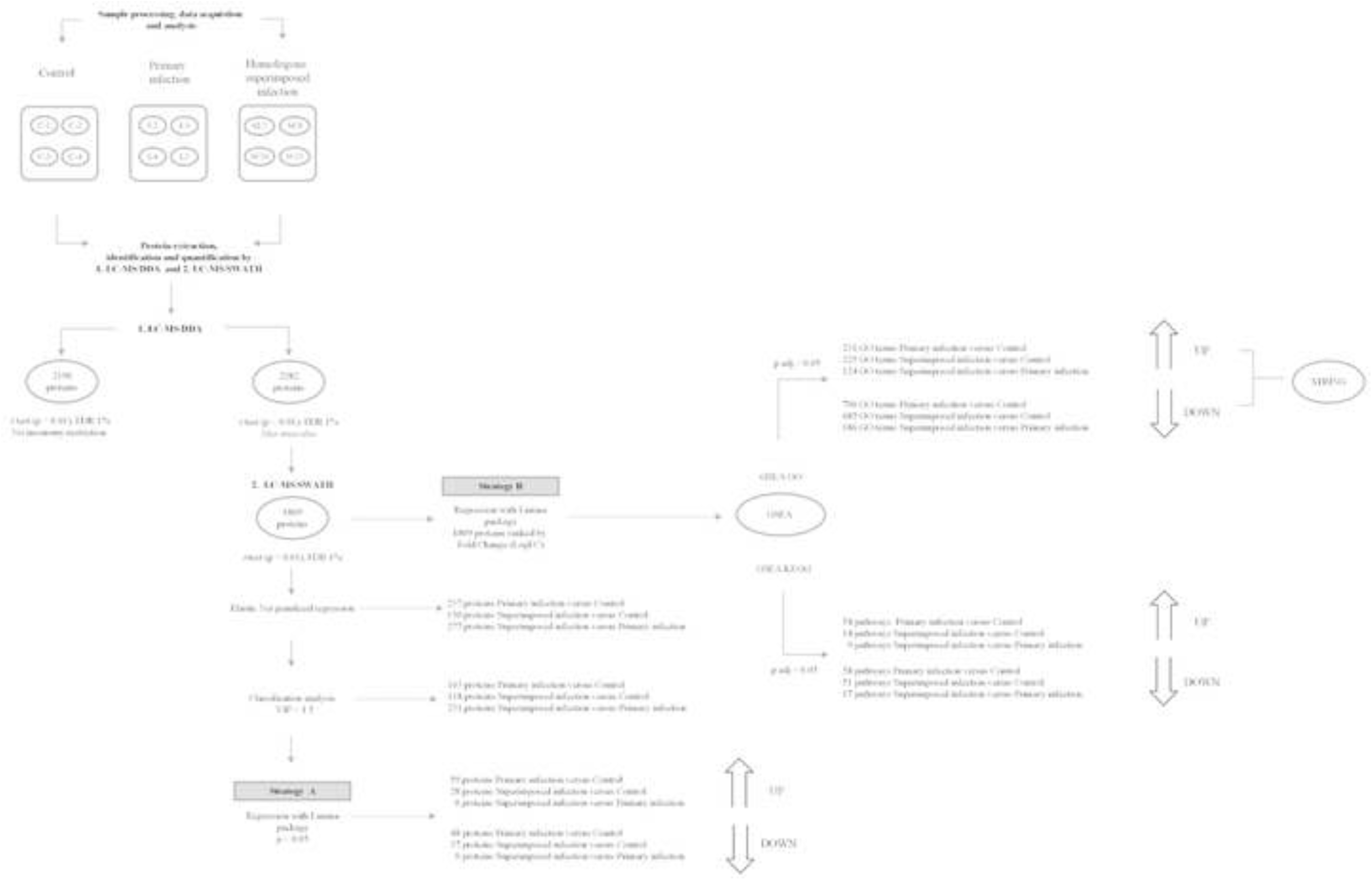
Experimental design and identification of differentially expressed proteins in mouse ileum infected with *Echinostoma caproni*. Four biological samples were included per experimental group: control (C1–C4), primary infection (I2–I5), and homologous superimposed infection (SI7–SI12). Following protein extraction, liquid chromatography–tandem mass spectrometry (LC–MS/MS) was performed for protein identification and quantification. A total of 1,809 proteins were identified and analyzed using two complementary strategies: (A) a regularized regression model based on Elastic Net followed by supervised classification using Variable Importance in Projection (VIP > 1.5); and (B) fold change (FC)-based regression analysis followed by bioinformatic functional enrichment analyses, including Gene Set Enrichment Analysis with Gene Ontology (GSEA–GO), Gene Set Enrichment Analysis with Kyoto Encyclopedia of Genes and Genomes pathways (GSEA–KEGG), and protein–protein interaction analysis using STRING.

### Functional analysis of omics data (GSEA: GO and KEGG)

Functional enrichment analysis using the GO database revealed that, when homologous superimposed infection was compared with the control group, a total of 910 enriched functional annotations were detected, of which 225 corresponded to upregulated proteins and 685 to downregulated proteins. Among the 10 most relevant functional annotations, activation of biological processes related to the modification of cellular components and cell walls of other organisms, digestion, and negative regulation of type I interferon-mediated signaling pathways was observed. In addition, a marked overactivation of a broad group of exopeptidases, metalloexopeptidase, metallopeptidase and peptidase families was detected (Fig 4, Table 1), many of which were localized to the extracellular region and space.

**Fig 4.**
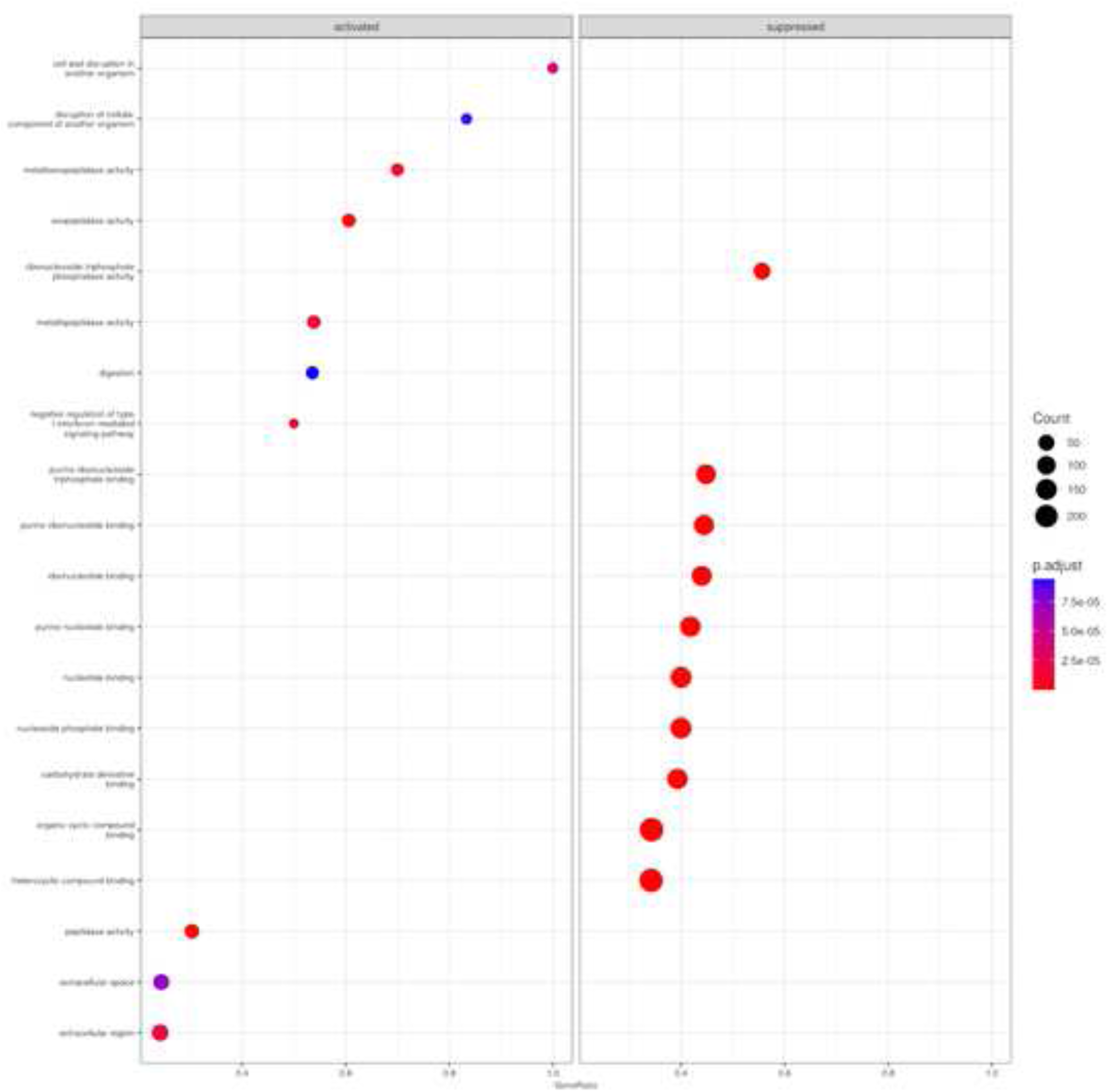
Principal GO terms using Gene Set Enrichment Analysis during superimposed *Echinostoma caproni* infection versus controls. Top 10 most representative GO terms associated with overexpressed (A) and underexpressed (B) proteins are shown. The size of the dots indicates the number of proteins mapped to each GO term, and the color represents the adjusted p-value.

**Table 1.** GO-based functional enrichment analysis using Gene Set Enrichment Analysis during superimposed *Echinostoma caproni* infection versus controls.

| GO ID | Expression | Description | ES | NES | p-value | Proteins |
| --- | --- | --- | --- | --- | --- | --- |
| MF GO:0008238 | ↑ | exopeptidase activity | 0.674 | 2.47 | 2.16<sup>-07</sup> | Anpep/Cj |
| MF GO:0008235 | ↑ | metalloexopeptidase activity | 0.740 | 2.42 | 1.00<sup>-05</sup> | Anpep/Cj |
| MF GO:0008237 | ↑ | metallopeptidase activity | 0.587 | 2.21 | 1.33<sup>-05</sup> | Anpep/M |
| BP GO:0007586 | ↑ | digestion | 0.618 | 2.14 | 9.42<sup>-05</sup> | Ctrb1/Fal |
| MF GO:0008233 | ↑ | peptidase activity | 0.459 | 2.11 | 9.57<sup>-07</sup> | Ctrb1/An |
| BP GO:0140975 | ↑ | disruption of cellular component of another organism | 0.974 | 2.02 | 9.00<sup>-05</sup> | Reg2/Reg |
| BP GO:0060339 | ↑ | negative regulation of type I interferon-mediated signaling pathway | 0.983 | 1.88 | 9.64<sup>-06</sup> | Isg15/Oa |
| BP GO:0044278 | ↑ | cell wall disruption in another organism | 0.914 | 1.86 | 4.56<sup>-05</sup> | Reg2/Reg |
| CC GO:0005576 | ↑ | extracellular region | 0.311 | 1.58 | 1.62<sup>-05</sup> | Reg2/Isg |
| CC GO:0005615 | ↑ | extracellular space | 0.312 | 1.56 | 7.44<sup>-05</sup> | Reg2/Am |
| MF GO:0035639 | ↓ | purine ribonucleoside triphosphate binding | -0.553 | -2.20 | 1.00<sup>-10</sup> | Psmc6/C |
| MF GO:0032555 | ↓ | purine ribonucleotide binding | -0.551 | -2.20 | 1.00<sup>-10</sup> | Psmc6/C |
| MF GO:0017111 | ↓ | ribonucleoside triphosphate phosphatase activity | -0.592 | -2.20 | 1.00<sup>-10</sup> | Arf4/Ar1 |
| MF GO:0032553 | ↓ | ribonucleotide binding | -0.547 | -2.18 | 1.00<sup>-10</sup> | Psmc6/C |
| MF GO:0017076 | ↓ | purine nucleotide binding | -0.523 | -2.11 | 1.00<sup>-10</sup> | Psmc6/C |
| MF GO:0000166 | ↓ | nucleotide binding | -0.499 | -2.02 | 1.00<sup>-10</sup> | Psmc6/C |
| MF GO:1901265 | ↓ | nucleoside phosphate binding | -0.499 | -2.02 | 1.00<sup>-10</sup> | Psmc6/C |
| MF GO:0097367 | ↓ | carbohydrate derivative binding | -0.462 | -1.86 | 1.00<sup>-10</sup> | Psmc6/C |
| MF GO:0097159 | ↓ | organic cyclic compound binding | -0.441 | -1.80 | 1.00<sup>-10</sup> | Rbm39/R |
| MF GO:1901363 | ↓ | heterocyclic compound binding | -0.438 | -1.98 | 1.00<sup>-10</sup> | Rbm39/R |

At the metabolic level, enrichment of proteins with mitochondrial activity persisted, particularly those associated with ATP synthesis coupled to the electron transport chain and with deoxyribonucleotide metabolism, together with activation of processes mediated by antimicrobial peptides. In addition, although not among the top enriched terms, overexpression of functional annotations related to hypoxia response and epithelial remodeling was also observed (S1 Table). In contrast, a sustained decrease in the expression of proteins involved in binding to purine ribonucleoside triphosphates, ribonucleotides, carbohydrate derivatives, cyclic organic compounds and heterocyclic organic compounds was detected (Fig 4, Table 1).

When homologous superimposed infection was compared with primary infection, 310 coordinately enriched functional annotations were identified. Upregulated proteins clustered into 124 functional terms, with the 10 leading GO terms associated with keratinization, intermediate filament cytoskeleton organization and keratinocyte differentiation. These proteins were predominantly localized to intermediate and keratin filaments, as well as to cornified envelope. Enriched terms related to proteins localized in the extracellular region and space were also identified (Fig 5, Table 2). Moreover, although not among the most highly enriched terms, overexpression of functional annotations associated with cellular response to oxygen levels, hypoxia and activation of peroxisomal matrix was noted. Likewise, results showed an enrichment of marker of intestinal epithelial differentiation and intestinal vascular permeability (S2 Table). Downregulated proteins were associated with enrichment of 186 functional annotations, with the top 10 GO terms indicating negative regulation of proteins involved in mitochondrial structure-mediated cellular metabolism, particularly those related to the mitochondrial respirasome, inner mitochondrial membrane and mitochondrial envelope, as well as the respiratory chain complex and aerobic respiration (Fig 5, Table 2).

**Fig 5.**
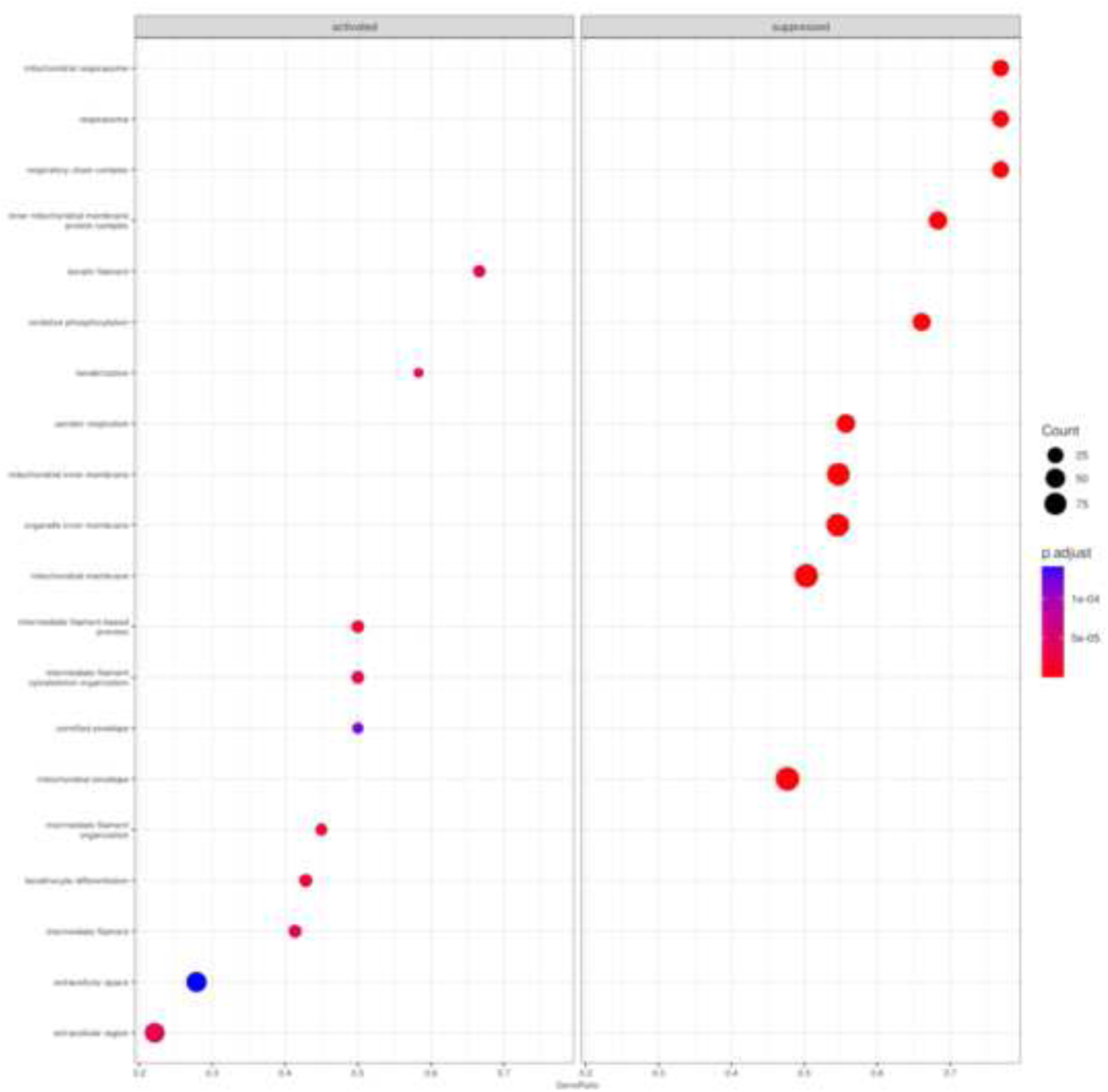
Principal Go terms using Gene Set Enrichment Analysis during superimposed versus primary *Echinostoma caproni* infections. Top 10 most representative GO terms associated with overexpressed (A) and underexpressed (B) proteins are shown. The size of the dots indicates the number of proteins mapped to each GO term, and the color represents the adjusted p-value.

**Table 2.** GO-based functional enrichment analysis using Gene Set Enrichment Analysis during superimposed versus primary *Echinostoma caproni* infections.

| GO ID | Expression | Description | ES | NES | p-value | Proteins |
| --- | --- | --- | --- | --- | --- | --- |
| BP GO:0045109 | ↑ | intermediate filament organization | 0.761 | 2.19 | 1.55<sup>-05</sup> | Krt16/Kr |
| BP GO:0045103 | ↑ | intermediate filament-based process | 0.739 | 2.18 | 2.28<sup>-05</sup> | Krt16/Kr |
| BP GO:0045104 | ↑ | intermediate filament cytoskeleton organization | 0.739 | 2.18 | 2.28<sup>-05</sup> | Krt16/Kr |
| BP GO:0031424 | ↑ | keratinization | 0.851 | 2.18 | 3.40<sup>-05</sup> | Krt16/Kr |
| CC GO:0045095 | ↑ | keratin filament | 0.809 | 2.16 | 4.10<sup>-05</sup> | Krt16/Kr |
| BP GO:0030216 | ↑ | keratinocyte differentiation | 0.699 | 2.15 | 1.69<sup>-05</sup> | Krt16/Kr |
| CC GO:0005882 | ↑ | intermediate filament | 0.681 | 2.11 | 3.56<sup>-05</sup> | Krt16/Kr |
| CC GO:0001533 | ↑ | cornified envelope | 0.770 | 2.10 | 0.0001 | Krt16/Kr |
| CC GO:0005576 | ↑ | extracellular region | 0.344 | 1.57 | 3.73<sup>-05</sup> | Mcpt1/M |
| CC GO:0005615 | ↑ | extracellular space | 0.353 | 1.57 | 0.0001 | Mcpt1/M |
| CC GO:0005746 | ↓ | mitochondrial respirasome | -0.710 | -2.13 | 1.06<sup>-06</sup> | Ndufb10/ |
| CC GO:0070469 | ↓ | respirasome | -0.710 | -2.13 | 1.06<sup>-06</sup> | Ndufb10/ |
| CC GO:0098803 | ↓ | respiratory chain complex | -0.710 | -2.13 | 1.06<sup>-06</sup> | Ndufb10/ |
| CC GO:0098800 | ↓ | mitochondrial inner membrane protein complex | -0.650 | -2.11 | 1.98<sup>-07</sup> | Atp5a1/A |
| BP GO:0006119 | ↓ | oxidative phosphorylation | -0.644 | -2.08 | 6.07<sup>-07</sup> | Atp5a1/A |
| CC GO:0019866 | ↓ | organelle inner membrane | -0.557 | -2.05 | 2.98<sup>-09</sup> | Cnp/Ak2 |
| CC GO:0005743 | ↓ | mitochondrial inner membrane | -0.554 | -2.04 | 4.36<sup>-09</sup> | Cnp/Ak2 |
| BP GO:0009060 | ↓ | aerobic respiration | -0.588 | -1.97 | 2.29<sup>-06</sup> | Atp5e/Dl |
| CC GO:0031966 | ↓ | mitochondrial membrane | -0.489 | -1.83 | 1.21<sup>-06</sup> | Atp5e/Hk |
| CC GO:0005740 | ↓ | mitochondrial envelope | -0.481 | -1.81 | 1.28<sup>-06</sup> | Hk1/Cyb |

↑ Overexpressed terms; ↓ Underexpressed term; ES, Enrichment Score; GO, Gene Ontology;
BP, biological processes; CC, cellular components; NES, Normalized Enrichment Score.

KEGG pathway analysis comparing homologous superimposed infection with the control group identified 14 activated and 51 inhibited pathways. Activated pathways included protein and carbohydrate digestion and absorption, NOD-like receptor signaling, and peroxisome proliferator-activated receptor (PPAR) signaling. In contrast, inhibited pathways encompassed motor proteins, phagosome, mitogen-activated protein kinases (MAPK) signaling, Fc gamma receptor-mediated phagocytosis, Ras signaling, endocytosis, vascular endothelial growth factor (VEGF) signaling, gap junction and tight junction integrity, cytoskeleton regulation, platelet activation, and phospholipase D and Rap1 signaling pathways (Fig 6).

**Fig 6.**
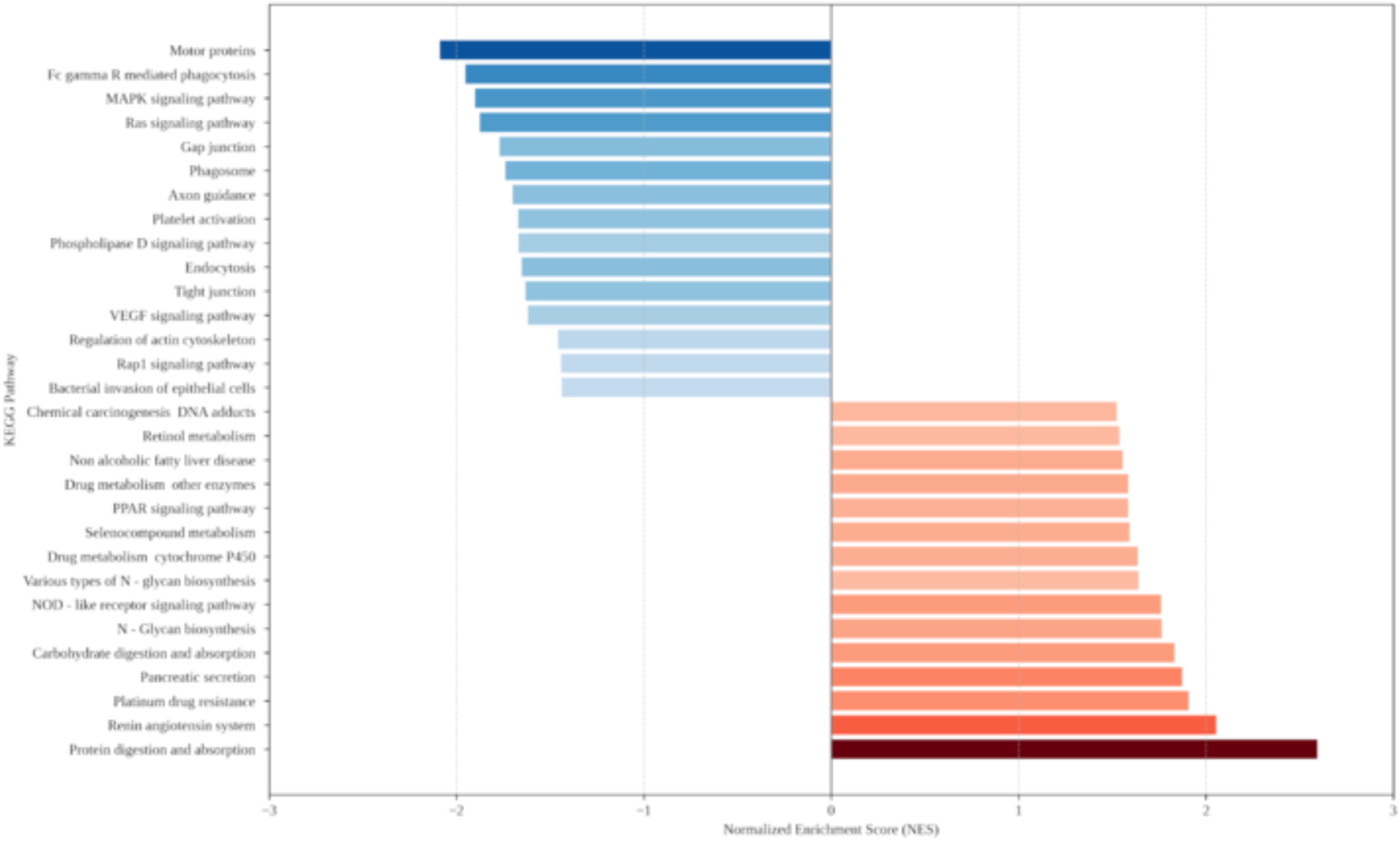
Principal pathways in ileum of mice during superimposed *Echinostoma caproni* infection versus controls. The graph shows the signaling pathways and metabolic processes enriched using Kyoto Encyclopedia of Genes and Genomes (KEGG), sorted by their Normalized Enrichment Score (NES) and indicating overactivated (red) and suppressed (blue) pathways.

Comparison of homologous superimposed infection with primary infection revealed 9 activated and 17 suppressed pathways, including activation of glycerophospholipid metabolism, alpha-linolenic acid metabolism and lysosomal components (Fig 7), together with suppression of oxidative phosphorylation, chemical carcinogenesis by reactive oxygen species (ROS), the cGMP–PKG signaling pathway, motor proteins, fatty acid biosynthesis and neurodegeneration-related pathways (S2 Table).

**Fig 7.**
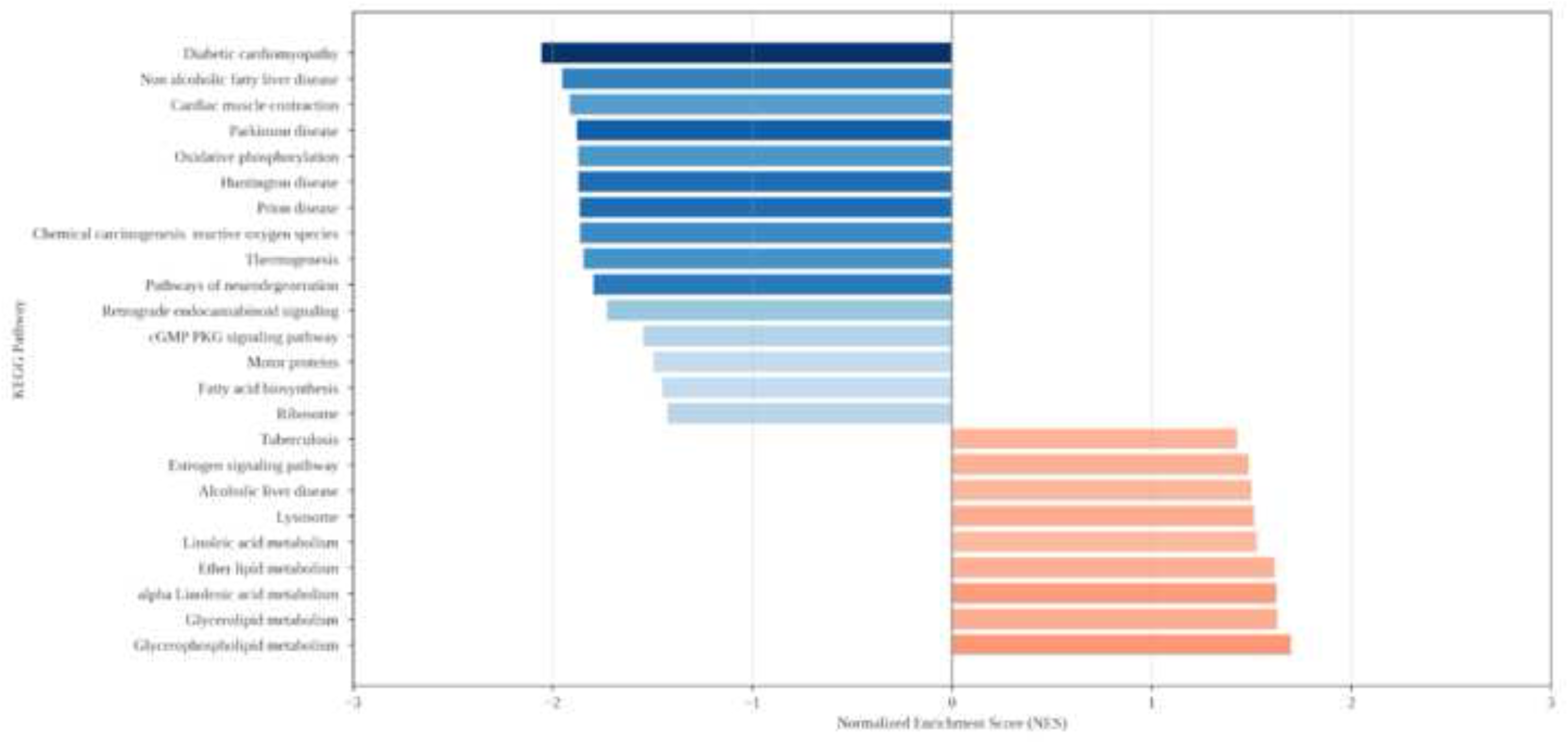
Principal pathways in ileum of mice during superimposed and primary *Echinostoma caproni* infections. The graph shows the signaling pathways and metabolic processes enriched using Kyoto Encyclopedia of Genes and Genomes (KEGG), sorted by their Normalized Enrichment Score (NES) and indicating overactivated (red) and suppressed (blue) pathways.

### Protein–protein interaction analysis (STRING)

STRING-based analysis of proteins grouped within the top 20 GO-based GSEA annotations revealed interaction patterns consistent with the functional enrichment results. In the comparison between homologous superimposed infection and the control group, upregulated proteins clustered into three major groups: proteins with exopeptidase and metalloprotease activity (cluster 1), proteins involved in lipid digestion and cell wall degradation of other organisms (cluster 2), and proteins associated with antigen cross-presentation (cluster 3) (Fig 8). Downregulated proteins formed clusters related to cellular amide metabolism (cluster 1), GTP binding (cluster 2), and antiparasitic responses against helminths (cluster 3) (Fig 9).

**Fig 8.**
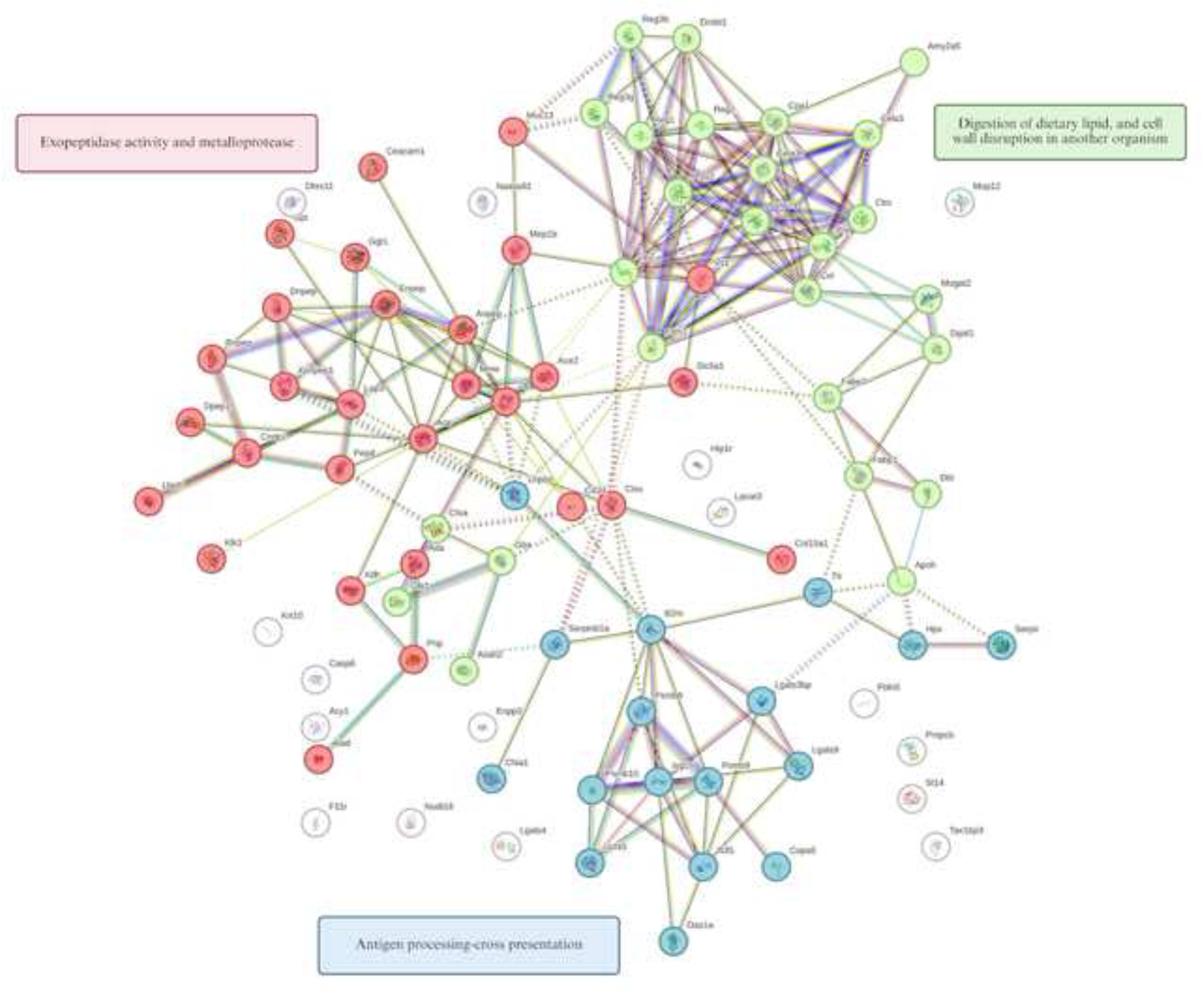
Protein-protein interaction from overexpressed proteins during superimposed *Echinostoma caproni* infection versus controls. Proteins were grouped into the top 10 GO terms using GSEA analysis. Hierarchical grouping by STRING of overexpressed proteins into three clusters is observed. Node size represents the relative centrality of each protein in the network and the color reflects the degree of connectivity or interaction density within the group. Lines indicate the type of interaction evidence supporting the connection between nodes (experimental data, coexpression, or curated databases).

**Fig 9.**
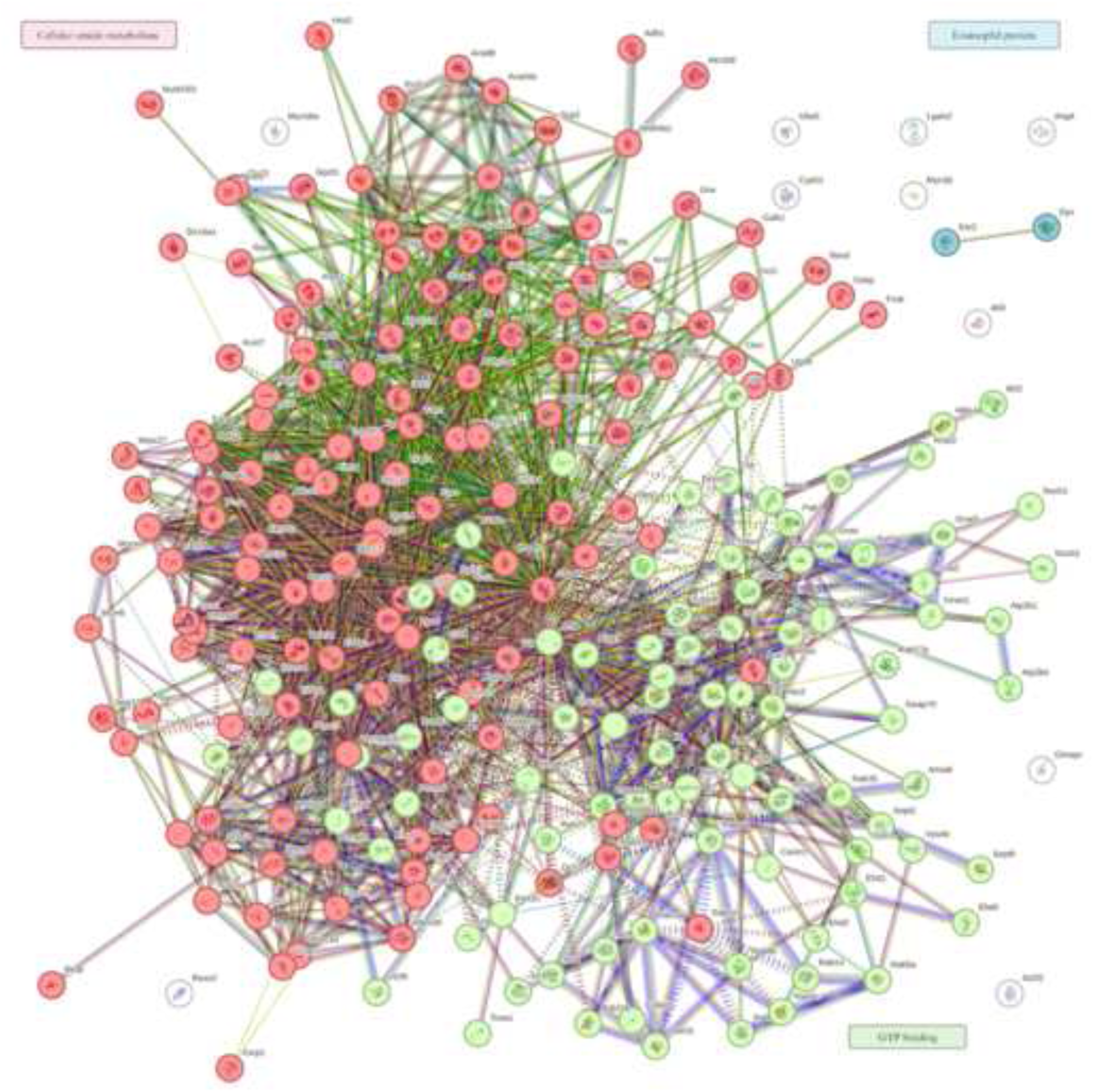
Protein-protein interaction from underexpressed proteins during superimposed *Echinostoma caproni* infection versus controls. Proteins were grouped into the top 10 GO terms using GSEA analysis. Hierarchical grouping by STRING of underexpressed proteins into three clusters is observed. Node size represents the relative centrality of each protein in the network and the color reflects the degree of connectivity or interaction density within the group. Lines indicate the type of interaction evidence supporting the connection between nodes (experimental data, coexpression, or curated databases).

Finally, comparison between homologous superimposed infection and primary infection revealed 6 main clusters grouping proteins involved in intermediate filament cytoskeleton organization, TIL (trypsin inhibitor-like cysteine-rich) domains and oligosaccharide binding, lipid digestion and cell wall degradation of other organisms (cluster 1); proteins forming extracellular matrix complexes with collagen (cluster 2); proteins associated with the IgE receptor (cluster 3); proteins involved in the pentose phosphate pathway and glycolysis (cluster 4); proteins associated with pyrimidine ribonucleotide metabolism (cluster 5); and proteins with antimicrobial activity (cluster 6) (Fig 10). Downregulation was primarily centered on mitochondrial proteins essential for oxidative phosphorylation, electron transport, ATP synthesis and heat production (cluster 1), while additional minor clusters included proteins involved in arginine biosynthesis (cluster 2) and nitrite reductase and NADH-related activity (cluster 3) (Fig 11).

**Fig 10.**
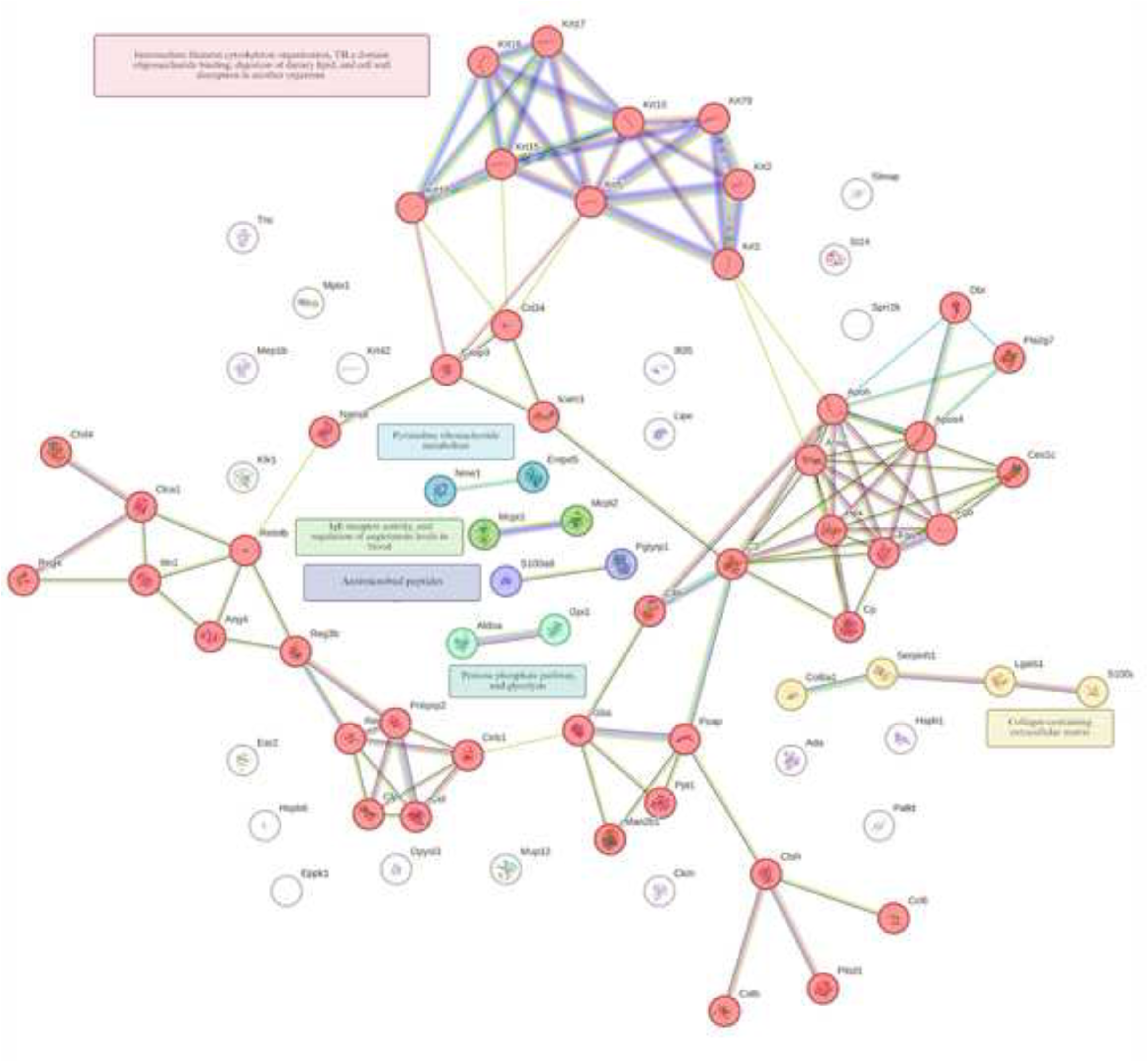
Protein-protein interaction from overexpressed proteins during superimposed and primary *Echinostoma caproni* infections. Proteins were grouped into the top 10 GO terms using GSEA analysis. Hierarchical grouping by STRING of overexpressed proteins into 6 clusters is observed. Node size represents the relative centrality of each protein in the network and the color reflects the degree of connectivity or interaction density within the group. Lines indicate the type of interaction evidence supporting the connection between nodes (experimental data, coexpression, or curated databases).

**Fig 11.**
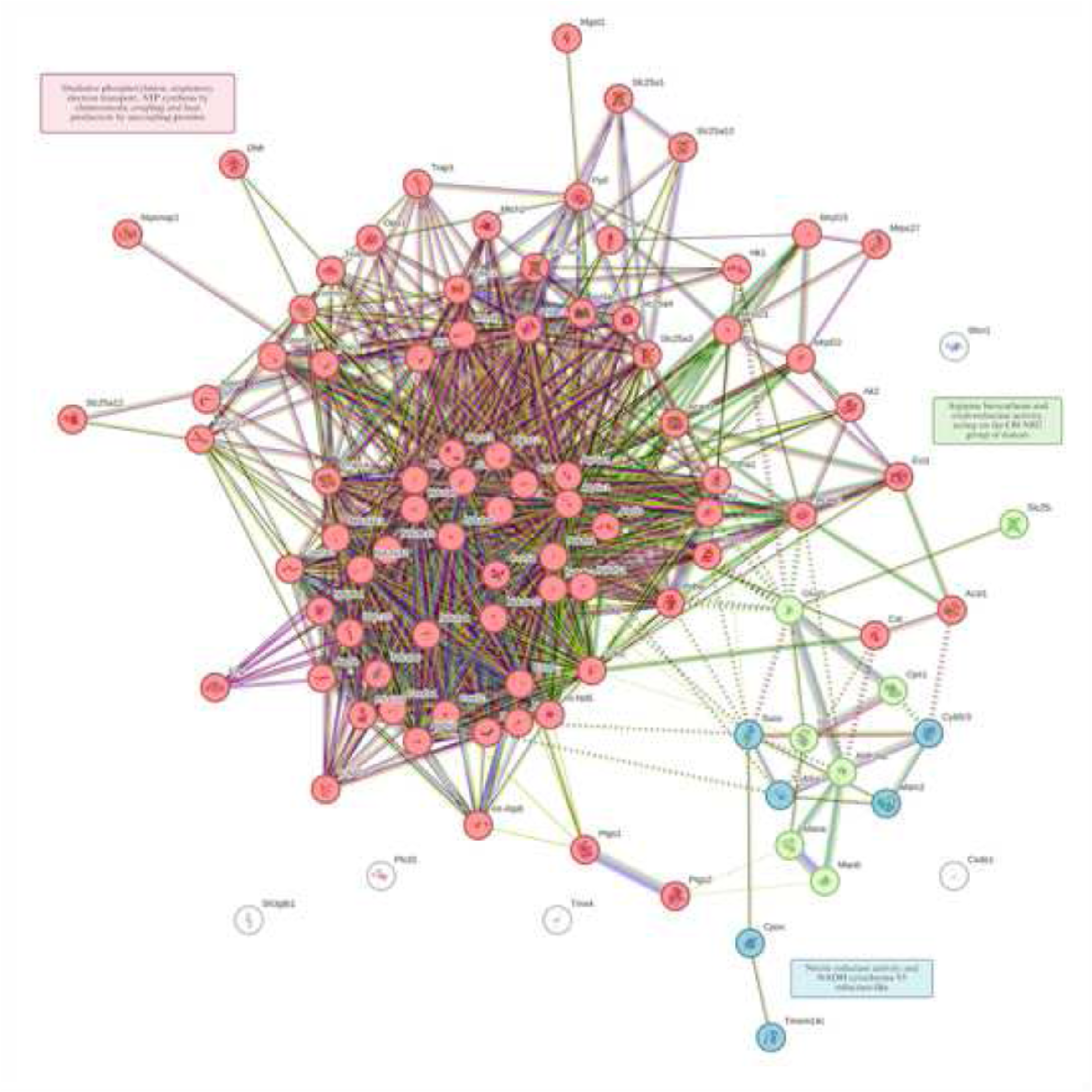
Protein-protein interaction from underexpressed proteins during superimposed and primary *Echinostoma caproni* infections. Proteins were grouped into the top 10 GO terms using GSEA analysis. Hierarchical grouping by STRING of underexpressed proteins into three clusters is observed. Node size represents the relative centrality of each protein in the network and the color reflects the degree of connectivity or interaction density within the group. Lines indicate the type of interaction evidence supporting the connection between nodes (experimental data, coexpression, or curated databases).

## Discussion

This study explored proteomic changes in intestinal epithelial cells associated with the loss or partial maintenance of intestinal homeostasis during the homologous superimposed infection with *E. caproni*. Homologous superimposed infections showed a markedly reduced infection rate compared with primary infection, confirming the development of partial resistance to repeated parasite exposure. This resistance has been largely attributed to the immunoregulatory role of IL-25 in the intestinal epithelium, as its increased activity correlates with reduced parasite burden [10]. However, whether repeated exposure induces adaptive or compensatory mechanisms at the proteomic level had not been previously addressed. Our study shows that despite univariate analysis did not detect any single protein with significant differential expression (after FDR correction) between the primary and superimposed infection groups, integrative multivariate modeling and gene set enrichment at the pathway level demonstrated a coordinated shift in epithelial cell function, supporting partial recovery of intestinal homeostasis.

Primary *E. caproni* infection has been shown to profoundly disrupt intestinal epithelial metabolism, differentiation, and cell death processes, promoting a persistent inflammatory microenvironment sustained by alternative oxidative metabolic pathways [7,18–21]. This chronic state has been linked to deficient IL-25 production, which may contribute to the host’s inability to eliminate the parasite [19].

Comparative proteomic analysis revealed significant differences between infected and control mice but no major individual protein changes between primary infection and homologous superimposed infection, consistent with previous findings [20]. Nonetheless, pathway-level analyses revealed coordinated functional reprogramming across metabolic, signaling, and tissue repair pathways, supporting the concept that global pathway modulation rather than single-protein changes underlies the divergent outcomes of primary infection and homologous superimposed infection [5].

Primary infection was characterized by enrichment of ROS-mediated chemical carcinogenesis and suppression of pathways involved in oxygen transport, activation of the hypoxia-inducible factor (HIF-1), VEGF, nitric oxide (NO) metabolism, and canonical Wnt signaling [18]. In this context, several studies have shown that excessive accumulation of reactive oxygen species (ROS) profoundly disrupts cellular homeostasis [22]. However, cells exposed to these conditions can activate compensatory signaling pathways aimed at restoring redox imbalance [23–25]. Surprisingly, our proteomic data reveal a downregulation of cytoprotective and antioxidant proteins such as cytoglobin (Cybg) and hemoglobin beta subunits (Hbb-b1, Hbb-b2), pointing to impaired redox homeostasis and reduced intracellular oxygen availability, thereby favoring a hypoxic epithelial microenvironment. Although ROS can stabilize HIF-1α and promote VEGF-dependent angiogenesis [26–28], our data suggest suppression of these axes, accompanied by reduced glycolytic enzyme activity (Pgk1, Aldoa, Ldhb) and diminished VEGF signaling linked to Cdc42 and Mapk2. Together with reduced expression of NO-related proteins (Icam1, Itgb2, Ddah1, Ass1), these findings support impaired angiogenesis, revascularization, and tissue repair during primary infection.

Oxidative stress has also been reported to activate the Wnt/β-catenin pathway [29,30]; however, primary *E. caproni* infection showed downregulation of multiple Wnt-associated proteins, including Jup and Ctnnd1. Given that IL-25 treatment restores Jup expression and supports epithelial integrity [8], these findings support a model in which oxidative stress combined with IL-25 deficiency interferes with regenerative signaling, leading to sustained epithelial damage and loss of homeostasis.

In contrast, homologous superimposed infections exhibited a functional profile consistent with partial restoration of intestinal homeostasis. Enriched pathways included lysosomal activity, peroxisomal matrix, long-chain fatty acid metabolism, PPAR signaling, and hypoxia-adapted respiration, alongside reduced mitochondrial oxidative phosphorylation and ROS-mediated carcinogenesis. Increased expression of lysosomal enzymes (Ctsh, Ppt1, Man2b1, Psap, Gba) suggests enhanced mitophagy and lipid turnover, limiting mitochondrial ROS accumulation and promoting cellular homeostasis [31,32].

Activation of peroxisomal matrix and PPAR signaling was associated with upregulation of Fabp1, Acox3, Nudt7, and Abcd3, supporting a shift toward lipid-based energy metabolism and redox balance [33,34]. Given that IL-25 promotes the mitochondrial respiratory capacity through lipid utilization, PPAR activation and via M2 macrophage polarization, these pathways may represent adaptive mechanisms facilitating the intestinal homeostasis during homologous superimposed infection modulated for IL-25 [7,8,10].

Homologous superimposed infections also showed enrichment of epithelial differentiation and tissue remodeling processes, evidenced by increased expression of multiple keratins (Krt1, Krt2, Krt5, Krt10, Krt15–18, Krt42, Krt79) and proteins involved in cytoskeletal organization and barrier integrity intestinal [35–37]. Concurrent upregulation of proteins involved in the modulation of intestinal vascular permeability (Tjp1, Fermt2, Ceacam1, Ddah1) suggests reinforcement of epithelial barrier function, which may contribute to reduced permeability and tissue damage [38,39].

Additionally, homologous superimposed infection induced a qualitative shift in antimicrobial peptide (AMP) expression, including Ang4, Itln1a, Reg2, Reg3b, Reg4, Retnlb, S100a9, and Pglyrp1. Unlike the pro-inflammatory AMP profile observed during primary infection, this repertoire is consistent with a more regulated epithelial response, potentially limiting dysbiosis while enhancing anti-helminth defense. Given that Ang4 expression is linked to IL-25/IL-13 signaling and parasite expulsion in other helminth models [40–42], these changes may contribute to resistance upon repeated exposure. Finally, several studies have shown that interaction between IL-25 and its receptor activates multiple signaling pathways involved in self-renewal, cell survival and apoptosis [43–46]. In this context, STRING analysis revealed interactions among proteins associated with IgE receptor activation, notably mast cell protease isoforms 1 (Mcpt1) and 2 (Mcpt2). Importantly, IL-25 is essential for inducing IgE class switching, thereby enhancing recognition of helminth antigens and promoting antibody-dependent cellular cytotoxicity, in which eosinophils and mast cells release cytotoxic mediators that contribute to parasite expulsion [47]. Furthermore, our data revealed positive interactions among proteins associated with a collagen - rich extracellular matrix, including Col6a1, Serpinh1, Lgals1 y S100a11. Notably, IL-25 has been reported to upregulate profibrotic genes, such as type I and III collagens, thereby promoting fibroblast differentiation and collagen deposition [48]. Nevertheless, it is important to note that prolonged inflammation and sustained tissue remodeling increased the risk of cellular damage, underscoring the need for efficient clearance of defective and infected cells to prevent dysregulation and the potential emergence of malignant transformations in the host [21,49,50]. In this context, GSEA analysis has shown modulation of gasdermin-C2, which may indicate its potential contribution of regulated inflammatory cell death pathways. This mechanism could influence the removal of damaged or infected enterocytes and consequently limit tissular disruption during homologous superimposed infection. The gasdermin-C2 is a protein implicated in pyroptosis and host defense against intracellular and extracellular pathogens [51]. In experimental *Nippostrongylus brasiliensis* infection, gasdermin C upregulation and caspase-mediated cleavage have been associated with the lytic enterocyte death and parasite elimination through the release of antiparasitic factors into the intestinal lumen [52].

In conclusion, primary *E. caproni* infection induces profound intestinal epithelial dysfunction characterized by oxidative stress, suppression of hypoxia-responsive and regenerative pathways (HIF-1, VEGF, NO, Wnt), and impaired repair capacity in an IL-25-deficient environment [18]. In contrast, homologous superimposed infection triggers an adaptive functional reprogramming involving lysosomal and peroxisomal lipid metabolism, PPAR activation, cytoskeletal reorganization, barrier reinforcement, and a specialized AMP repertoire. Together, these responses support partial restoration of epithelial integrity and contribute to the intestinal homeostasis upon repeated parasite exposure. Although these conclusions are based on proteomic patterns and require functional validation, they support a coherent pathophysiological model integrating metabolism, signaling, tissue repair, and epithelial immunity (Fig 12), highlighting IL-25 as a potential central modulator of intestinal homeostasis during helminth infection.

**Fig 12.**
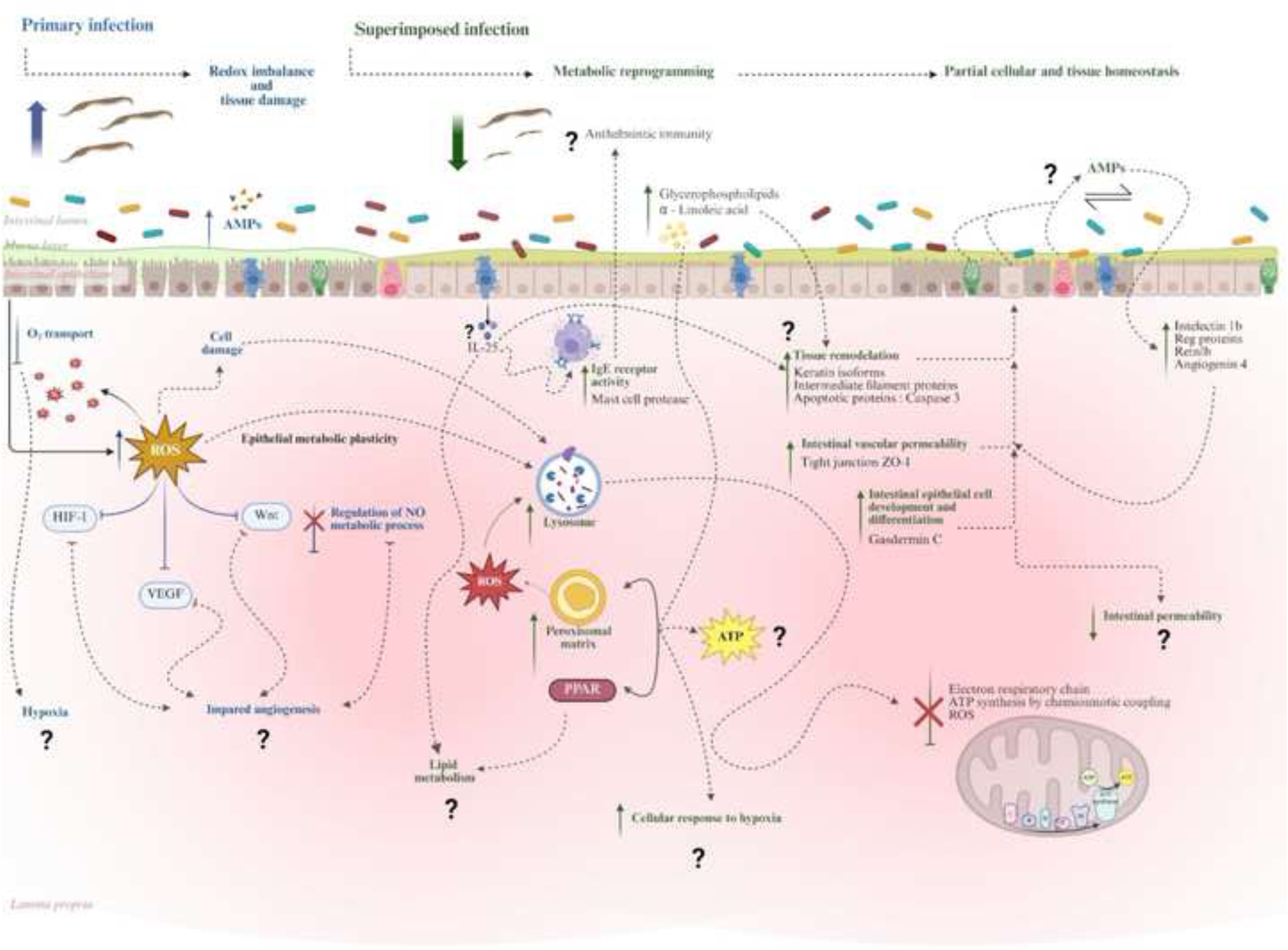
Schematic model of omics changes induced during primary and superimposed *Echinostoma caproni* infections. Primary *E. caproni* infection is associated with redox imbalance characterized by increased reactive oxygen species (ROS), altered oxygen transport, and suppression of hypoxia-inducible factor 1 (HIF-1), vascular endothelial growth factor (VEGF), and canonical Wnt signaling. These molecular changes may promote a hypoxic mucosal microenvironment and impaired angiogenesis in an IL-25-deficient context, contributing to loss of intestinal homeostasis. In contrast, homologous superimposed infection is characterized by metabolic reprogramming towards lipid metabolism, activation of peroxisome proliferator-activated receptor (PPAR)-dependent peroxisomal pathways, increased lysosomal activity, and modulation of ROS production and ATP synthesis. This response is accompanied by partial tissue remodeling, regulation of structural proteins, potential alterations in vascular and epithelial permeability, and modulation of antimicrobial peptide (AMP) expression. IL-25 may coordinate these responses by promoting epithelial repair, metabolic plasticity, and partial resistance to homologous superimposed infection, contributing to partial restoration of cellular and tissue homeostasis. Dashed lines and question marks indicate hypothetical or not fully elucidated mechanisms. Blue: primary infection; green: homologous superimposed infection. Figure created in BioRender Fiallos, E. (2026) https://BioRender.com/uhcp3ud.

## Data Availability Statement

All proteomic data have been deposited to the ProteomeXchange Consortium via the PRIDE partner repository with the Project accession: PXD076260. The full list of data can be found in the Supporting Information files.

## Acknowledgements

This work was supported by the project UV-INV_AE_4211753 from the University of Valencia (Spain). We thank to Animal production and experimentation Units of the Central Support Service for Experimental Research (SCSIE) of the Universitat de València for technical assistance. The proteomic analysis was performed at the Proteomics Facility of SCSIE, that belongs to ProteoRed, and statistical analyses at the Statistics and Omics Data Analysis Section of SCSIE. EF and PC acknowledge support from the ACIF-Grisolía fellowship and APOSTD/2024 postdoctoral program of the Generalitat Valenciana (Spain), respectively.

## Supporting information

S1 file. R script utilized for GSEA comparation between superimposed *Echinostoma caproni* infection and controls.

S2 file. R script utilized for GSEA comparation between superimposed and primary *Echinostoma caproni* infections.

S1 Table. GO terms identified by GSEA and KEGG during superimposed *Echinostoma caproni* infection versus controls.

S2 Table. GO terms identified by GSEA and KEGG during superimposed versus primary *Echinostoma caproni* infections.

